# PfD123 modulates K13-mediated survival and recovery after artemisinin exposure

**DOI:** 10.1101/2022.01.27.476788

**Authors:** Christopher Nötzel, Björn F. C. Kafsack

## Abstract

Recent advances in curbing the deadly toll of malaria have been threatened by the emergence of parasites resistant to the front-line antimalarial artemisinin. Resistance is mediated by point-mutations in the parasite protein Kelch13, but the mechanism of resistance is multi-factorial and only partially understood. Resistance-conferring Kelch13 mutations have been shown to lead to low-level activation of the parasite’s integrated stress response (ISR) which has a protective effect against artemisinin through an unclear mechanism. Furthermore, only a subpopulation of resistant parasites ever survives drug exposure, implying an underlying heterogeneity. By applying scRNAseq to the resistance-relevant early ring stage, we found expansion of a subpopulation in Kelch13 mutant parasites that is chiefly characterized by transcription of the putative positive translational regulator D123, while we conversely observed reduced D123 protein levels at the same stage. Analogous inverse changes in D123 expression are produced by experimental activation of the ISR, and genetically manipulating D123 expression modulates sensitivity to artemisinin, establishing it as a stress-responsive gene that contributes to artemisinin resistance in Kelch13-mutant malaria parasites.

## INTRODUCTION

Malaria remains a major global health and economic burden, with around 200 million infections causing 409,000 deaths annually (World Health Organization, 2020). Virtually all clinical symptoms of the disease are caused by asexual replicative growth of the unicellular protozoan parasite within red blood cells (Phillips et al., 2017), and this stage is the main target of all drugs commonly used for treatment, including the current front-line antimalarial artemisinin (Menard & Dondorp, 2017). Artemisinin and its semisynthetic derivatives (collectively referred to as ARTs) are activated via free heme released during hemoglobin digestion by the parasite, thereby confining its action to infected red blood cells (Wang et al., 2015). Activation involves heme-mediated cleavage of the molecule’s endoperoxide bond, leading to highly reactive ART intermediates that inflict wide-spread cytotoxic damage and lead to activation of the parasite’s integrated stress response (ISR) that is poorly reversible (Bridgford et al., 2018). Importantly, efficient translational shutdown at the time of artemisinin exposure, as well as eventually overcoming it after drug levels subside are crucial for the parasite’s ability to recover from artemisinin treatment (Zhang et al., 2017).

Resistance to ARTs emerged initially in Western Cambodia (Dondorp et al., 2009; Noedl et al., 2009), is now widespread across Southeast Asia (Imwong et al., 2020) and has recently been found in Africa as well (Balikagala et al., 2021). Mutations in the parasite protein PfKelch13 are the molecular determinant of ART-resistance (Ariey et al., 2014; Straimer et al., 2015). One major resistance mechanism is the reduced release of free heme in K13-mutant early rings due to decreased hemoglobin digestion, as K13 is involved in hemoglobin uptake at that stage and mutations in K13 lead to reduced K13 abundance (Birnbaum et al., 2020; Yang et al., 2019; Gnädig et al., 2020; Stokes et al., 2021). Another critical contributor to resistance is an increased base-line activation of the parasite’s stress response that was detected in Kelch13-mutant parasites in the field (Mok et al., 2015) and was also shown to be important for resistance in vitro (Cui et al., 2012; Dogovski et al., 2015; Rocamora et al., 2018). Specifically, K13-mutant rings show increased levels of eIF2α phosphorylation at the resistance-relevant early ring stage (Zhang et al., 2017). In addition, experimentally activating the ISR using the eIF2α phosphatase inhibitor Salubrinal prior to exposure to ARTs provides a protective effect (Zhang et al., 2017). However, how this increased ISR activation exactly promotes resistance to ARTs remains unclear. It is conceivable that specific downstream effectors of the ISR are changed in expression in K13-mutant early rings, but our knowledge about the *Plasmodium* ISR is limited. Furthermore, the observation that only a subpopulation of K13-mutant rings exposed to ARTs ever survive drug treatment implies that any underlying changes in gene expression are restricted to a subpopulation of parasites (Straimer et al., 2015), and the use of conventional bulk transcriptomics approaches has made a detailed characterization of this heterogeneity challenging.

Here we used scRNAseq to characterize both the transcriptional heterogeneity at the resistance-relevant 0-3 hpi ring stage in ART-sensitive and -resistant parasites, as well as the heterogenous transcriptional response of parasites to ART exposure. We identify the putative positive translational regulator D123 as a major stress-responsive gene that is differentially expressed in K13-mutant early rings and find that genetically modulating D123 expression levels leads to changes in ART-sensitivity. Taken together, this work underscores the importance of a modified integrated stress response in ART-resistance and identifies a specific downstream effector of the *Plasmodium* ISR that contributes to ART-resistance in K13-mutant parasites.

## RESULTS

### The putative translational regulator D123 is differentially expressed in Kelch13-mutant, artemisinin-resistant early ring stage parasites

We hypothesized that mutations in Kelch13 lead to changes in gene expression within a subpopulation of early ring stage parasites, and that these pre-existing differences promote resistance to Artemisinin. To uncover any potential baseline changes in the transcriptional heterogeneity in Kelch13 mutant parasites, we performed single-cell transcriptional profiling of ring stage parasites immediately following invasion (0-3 hpi) in asexual Dd2 strain parasites carrying either the PfKelch13 wildtype allele (K13^wt^) or the ART-resistance-conferring R539T allele (K13^R539T^, Straimer et al., 2015, Supplementary Fig. 1a) in three biological replicates.

Generating 0-3 hpi samples generally yielded cultures at around 2-5% parasitemia, but scRNAseq requires input samples with a maximized fraction of infected red blood cells (iRBCs) to avoid wasteful sequencing of uninfected red blood cells (uRBCs). Preparing such samples from ring stage cultures is challenging as they lack the paramagnetic hemozoin present in more mature intra-erythrocytic stages that helps separating those from uRBCs with the help of magnetic columns. An alternative would be flow-cytometry-based sorting of iRBCs, but here the lack of flexibility when relying on institutional core facilities, as well as high costs and a relatively long duration of sorting are of concern. Instead, we employed a recently published method that relies on differences in the lipid composition between iRBCs and uRBCs to selectively enrich for iRBCs using the bacterial virulence factor streptolysin-O (Brown et al., 2020; see methods). Using this simple and quick protocol, we were able to reproducibly yield <70% iRBC samples that we could use for scRNAseq (Supplementary Fig. 1c).

Despite the narrow time window profiled and the very low RNA content of early ring stages (Martin et al., 2005; Sims et al., 2009; Supplementary Fig. 1b), we were able to resolve different subpopulations within each sample, with cells separating into two major clusters (1 and 2) in all three replicates (Fig. 1a, Supplementary Fig. 1d-e), as well as one or two minor clusters that varied by replicate. Marker analysis found that cells in cluster 2 were characterized by high levels of the DnaJ chaperone Ring Infected Surface Antigen (RESA, PfDd2_010006000). RESA is one of the most highly expressed genes immediately following merozoite invasion (Otto et al., 2010), thus validating this cluster 2 as containing nascent ring stages. K13^wt^ and K13^R539T^ parasites contributed similarly to cluster 2 and we found no significant expression differences between these strains among cluster 2 cells. Intriguingly, cluster 1 was significantly enriched for Kelch13-mutant parasites across all three biological replicates (1.68 fold-enrichment, p = 0.009, Fig. 1c-d, Supplementary Table 1) and was marked by expression of a gene encoding an uncharacterized D123-domain protein (PfDd2_030027500, “PfD123” hereafter). D123 proteins are broadly conserved across eukaryotes (Burroughs et al., 2015) and facilitate assembly of eIF2 translational initiation complex by functioning as a chaperone for the eIF2 subunits eIF2α and eIF2γ, thereby positively regulating protein translation (Bieganowski et al., 2004; Perzlmaier et al., 2013; Panvert et al., 2015).

**Figure 1:**
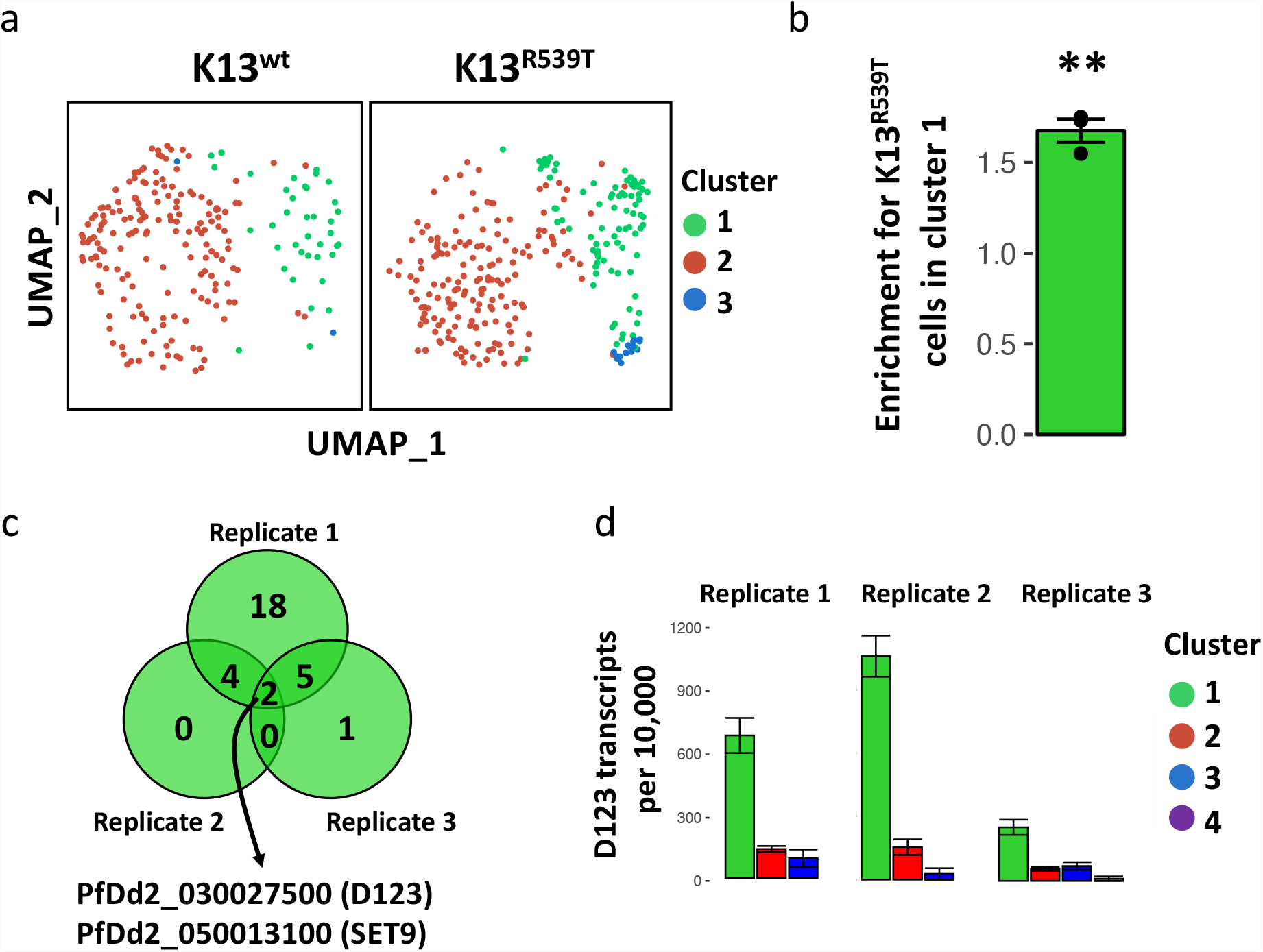
The putative translational regulator PfD123 is differentially expressed in K13-mutant, artemisinin-resistant early ring stage parasites. **a**, UMAP projection of replicate 1 of the 0-3 hpi scRNAseq experiment, stratified by parasite strain. **b**, enrichment of cluster 1 for K13^R539T^ parasites across the three 0-3 hpi scRNAseq replicates, n = 3, p = 0.009. **c**, venn diagram of markers identified for cluster 1 in the three 0-3 hpi replicates (Bonferroni corrected p-value < 0.05). **d**, average transcript levels of D123 detected per cell for each cluster and replicate. Error bars always represent sem, statistical significance in b was tested using two-sided, one-sample t-tests (mu = 1).

### Deconvoluting the heterogeneous transcriptional response to artemisinin

Kelch13-mutant parasites exhibit increased base-line activation of the integrated stress response (Zhang et al., 2017), which is known to lead to translational arrest in malaria parasites (Vembar et al., 2016). Since the regulation of protein translation is intimately connected to the ISR, we hypothesized that, as a putative translational regulator, expression of PfD123 might be stress-responsive and that increased levels of eIF2α phosphorylation previously described in early rings of Kelch13-mutants (Zhang et al., 2017) are responsible for the increased subpopulation of cells with high D123 transcript levels that we observed at that stage (Fig. 1). In order to test this, we performed scRNAseq of K13^wt^ and K13^R539T^ ring-stage parasites after exposure to Dihydroartemisinin (DHA) or a solvent control in a classic “ring survival assay” (RSA) setup (Witkowski et al., 2013): Tightly synchronous 0-3 hpi parasites were exposed to 700 nM DHA or DMSO for 6h, after which the drug was rigorously washed off and parasites were followed up for survival. Single-cell transcriptomics were performed at the wash-off time point (6-9 hpi), as well as 9 hours later (15-18 hpi) (n = 2, Supplementary Fig. 3a). Artemisinin itself leads to a potent activation of the ISR (Bridgford et al., 2018), making artemisinin-treated parasites a suitable model for stress-induced changes in gene expression. In addition, this experimental setup allowed us to better understand how parasites respond specifically to treatments with DHA, and whether surviving Kelch13-mutant parasites show any unique changes in gene expression after drug treatment, in addition to the pre-existing differential gene expression described in Figure 1.

When analyzed together, the single-cell transcriptomes (SCTs) of these RSA time-course experiments self-organized in a continuous arc along one side of the UMAP projection (clusters 2 – 5) with an additional, transcriptionally particularly distinct cluster 1 on the other side and one smaller cluster in-between (cluster 6), the latter of which was disregarded due to its small size (Fig. 2a). Visualizing the SCTs stratified by parasite strain, treatment and collection time point suggests that clusters 3 – 5 mainly represent healthy parasites progressing through the cell cycle, while clusters 1 and 2 are mainly composed of DHA-treated cells (Fig. 2b, Supplementary Fig. 3c). Enrichment analysis confirmed that clusters 1 and 2 are ART-response clusters (Fig. 2c), while correlation of pseudo-bulked SCTs to published bulk transcriptomics data validated that parasites progressively move through early intra-erythrocytic development from cluster 3 through cluster 5 (Supplementary Fig. 4).

**Figure 2:**
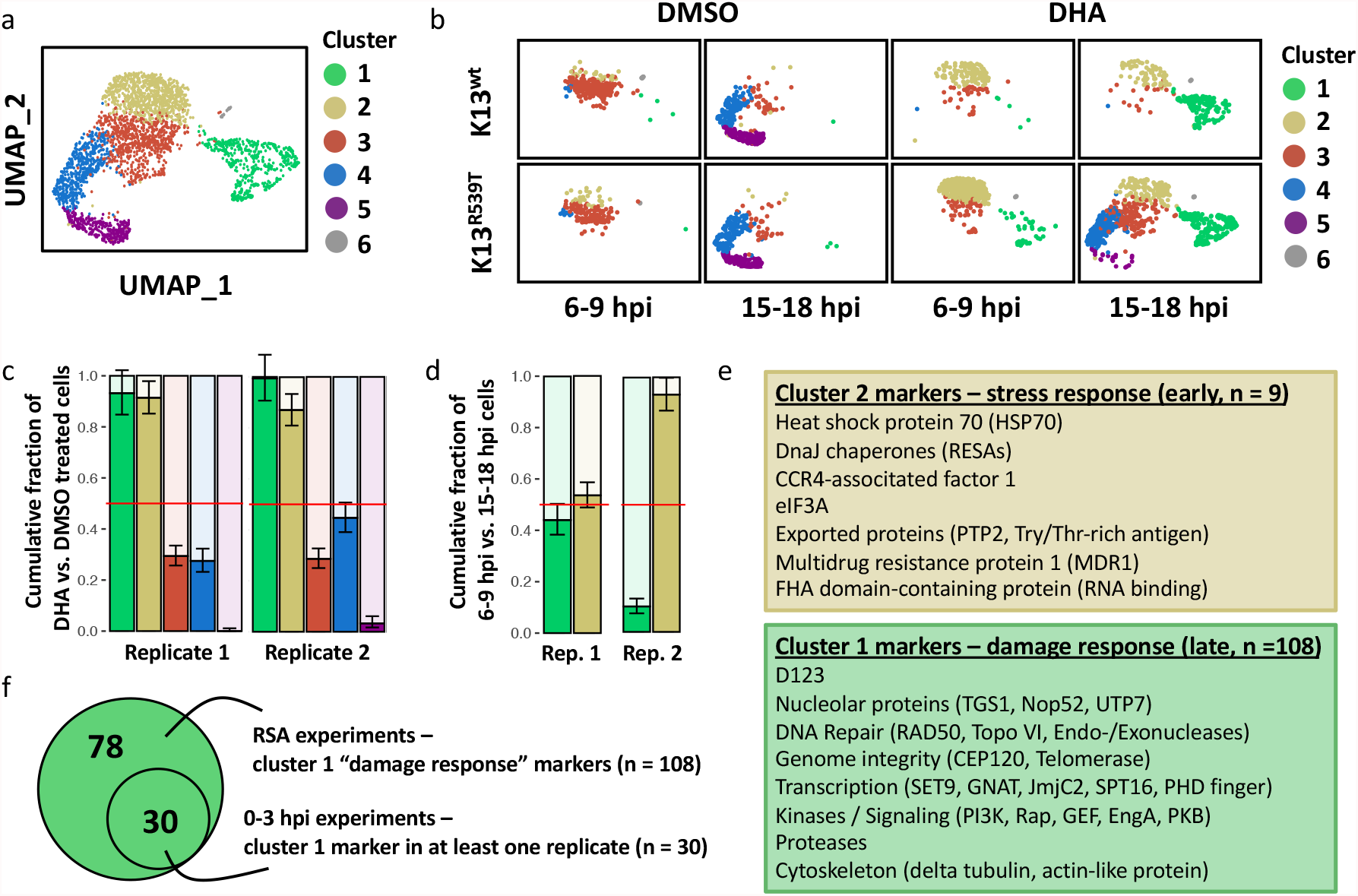
Deconvoluting the transcriptional response to artemisinin with single-cell resolution. **a**, UMAP projection of 6-9 hpi and 15-18 hpi SCTs from both DMSO and DHA treated K13^wt^ and K13^R539T^ parasites. **b**, same SCTs as in a, but stratified by strain, collection time point and treatment. **c**, for each cluster, enrichment for DHA vs. DMSO treated parasites was calculated, and the cumulative fraction plotted. Error bars represent 95% confidence intervals from poisson tests. Color code for clusters is the same as in b. **d**, for cluster 1 and 2, enrichment for 6-9 hpi vs. 15-18 hpi cells was calculated, and the cumulative fraction plotted. Error bars represent 95% confidence intervals from poisson tests. Color code for clusters is the same as in b. **e**, manually curated summary of marker genes for cluster 2 (top) and cluster 1 (bottom). Only genes that were significant markers for their respective cluster in both biological replicates were considered (p < 0.05, Bonferroni corrected p-value). **f**, venn diagram of cluster 1 markers of RSA experiments in this figure and cluster 1 markers of 0-3 hpi experiments (Fig. 1). DHA = Dihydroartemisinin.

Notably, the two ART-response clusters were clearly separated on the UMAP projection (Fig. 2a), indicating they both represent transcriptionally distinct stages of parasites responding to ART-induced cellular stress. Both clusters were each clearly enriched for one of the two collection time points, further suggesting that cluster 2 represents the immediate response to DHA (mostly 6-9 hpi cells, at the end of artemisinin treatment), while cluster 1 represents the later response (mostly 15-18 hpi cells collected 9h after washing off DHA) (Fig. 2d). The immediate stress response (cluster 2, Fig. 2e top panel) was characterized by chaperones and heat shock proteins, which help restricting protein damage, as well as by the Multidrug resistance protein (MDR1), which is a pump that decreases accumulation of antimalarials in the food vacuole and shows increased copy number in other drug-resistant parasites (Foote et al., 1989). eIF3A, the RNA-binding component of the translation initiation factor complex, as well as the RNA-regulatory proteins CCR4-associated factor 1 (CAF1) and FHA-domain containing protein were also expressed in early stress responsive cells. The later response to DHA (cluster 1, Fig. 2e bottom panel) showed a high number of reproducible marker genes (n = 108, compared to 9 markers for cluster 2) that were called with higher confidence based on adjusted p-values (Supplementary Table 2). These parasites were transcriptionally characterized by nucleolar proteins, DNA repair and genome integrity factors as well as many kinases and signaling proteins. Proteases, cytoskeletal proteins and regulators of transcription were also highly expressed. Overall, cluster 1 parasites appeared to mount a transcriptional response of recovery after the damage (“damage response”), indicative of an attempt to reenter the cell cycle after drug levels have subsided.

We were curious whether wildtype and Kelch13-mutant parasites respond differently to artemisinin or express different genes as they progress through early intra-erythrocytic development. However, when performing within-cluster marker analysis comparing gene expression between the two strains, no reproducible differences were found for any of the clusters in these experiments.

Kelch13-mutant parasites showed expected levels of survival after DHA-exposure in these RSA scRNAseq experiments (Supplementary Fig. 3a). Accordingly, we could identify a distinct subpopulation of K13-mutant parasites in the DHA-treated samples that was absent in Dd2 wildtype samples treated with DHA (Fig. 2b). This subpopulation appeared in both experiments, was largely restricted to the 15-18 hpi time-point (Supplementary Fig. 3c) and colocalized with untreated parasites moving through intra-erythrocytic development, altogether suggesting that these are parasites surviving and reentering the cell cycle after DHA-exposure. We were curious whether these cells showed any specific gene expression characterizing them as survivors. However, marker analysis comparing them to DMSO-treated parasites within the same global cluster showed no reproducible differences between the two experiments, indicating that at that point in time, K13-mutant survivors are transcriptionally largely comparable to healthy, asexually replicating parasites.

### Expression of PfD123 is a stress-responsive

Intriguingly, the marker lists of cluster 1 of the ring survival assay experiments (“damage response”) and of cluster 1 of the untreated 0-3 hpi experiment showed complete overlap (Fig. 2f), suggesting that those two subpopulations represent a similar transcriptional state. This supports our hypothesis that the parasites found in cluster 1 at 0-3 hpi (Fig. 1a) are characterized by a transcriptional response to stress, and that the expansion of this subpopulation in K13^R539T^ parasites (Fig. 1b) is likely a result of the pre-existing stress in those parasites. Importantly, this included D123 as one of the top markers of this damage response cluster, for which high transcript levels were detected in virtually every cell of this artemisinin-responsive subpopulation (Fig. 2e bottom panel, Fig. 3a-c).

**Figure 3:**
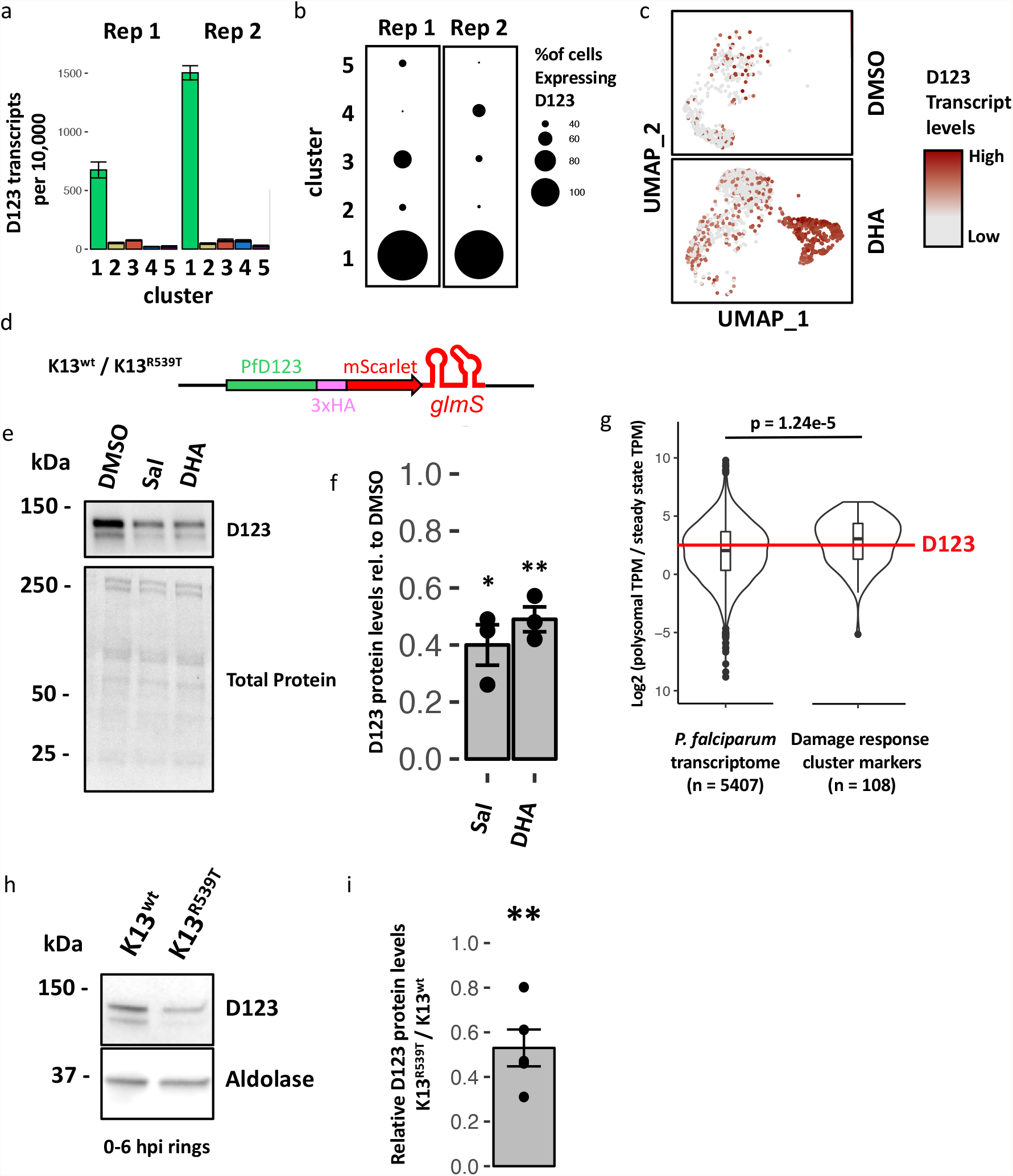
Expression of PfD123 is stress-responsive. **a**, Average transcript levels of D123 detected per cell after exposure to DHA for each cluster and replicate (RSA experiments Fig. 3.2). Error bars represent sem. **b**, Percent D123+ cells for each cluster, stratified by biological replicate, plotted in dot plots (RSA experiments Fig. 3.2). **c**, UMAP projection of replicate 2 of the RSA experiments (Fig. 3.2), stratified by treatment, with each cell colored based on the detected D123 transcript levels. **d**, schematic of K13^wt^-/ K13^R539T^ -D123-3xHA-mScarlet-glmS parasite lines generated by CRISPR/Cas9 gene editing **e**, representative immunoblot of K13^wt^-D123-3xHA-glmS ring stage parasites treated for 6h with 0.1% DMSO, 10 μM Salubrinal or 700 nM DHA. D123 was detected using an anti-HA antibody. Total protein on membrane was visualized using BioRad TGX stain-free gel technology. **f**, densitometry-based quantification of three replicates of the experiment shown in e, with D123 protein levels of Salubrinal or DHA treated parasites shown relative to those in DMSO treated parasites. D123 signal was first normalized to total protein signal. **g**, previously published polysome profiling data of ring stage parasites was reanalyzed (Bunnik et al., 2013). The log2 ratio of reads in the polysome-associated fraction vs. those in the steady-state mRNA fraction was calculated for each transcript as an estimate for its polysome occupancy. Violin diagrams and boxplots of the ratios were plotted for all *P. falciparum* transcript (left) or for the 108 transcripts of the RSA cluster 1 damage response marker genes (right). The red line indicates the ratio calculated for D123. Statistical significance was tested using a two-sided, unpaired t-test. **h**, representative immunoblot of 0-6 hpi K13^wt^-/ K13^R539T^ -D123-3xHA-mScarlet-glmS ring stage parasites. PfAldolase expression was used as a loading control. D123 was detected using an anti-HA antibody. **i**, densitometry-based quantification of D123 protein levels in 0-6 hpi rings, plotted as the ratio of Aldolase-normalized expression of D123 in K13^R539T^ -D123-3xHA-mScarlet-glmS over K13^wt^-D123-3xHA-mScarlet-glmS parasites, n = 5, p = 0.005. Representative replicate shown in h. Error bars always represent sem, statistical significance in f and i was tested using two-sided, one-sample t-tests (mu = 1, * = p < 0.05, ** = p < 0.01). Sal = Salubrinal, DHA = Dihydroartemisinin.

In order to track protein expression of D123, we generated the transgenic parasite lines K13^wt^-D123-3xHA-glmS and K13^R539T^-D123-3xHA-glmS, where the endogenous D123 locus is epitope-tagged with a 3xHA-mScarlet fusion followed by the glmS ribozyme sequence after the stop codon in order to allow for conditional knockdown (Prommana et al., 2013; Fig. 3d, Supplementary Fig. 2). We used this tool to test the effect of ISR activation on D123 protein levels by performing immunoblots after treating K13^wt^-D123-3xHA-glmS ring stage parasites for 6h with either 700 nM DHA or 10 μM of the eIF2α phosphatase inhibitor Salubrinal (Boyce et al., 2005), which both have been shown to robustly inflate eIF2α phosphorylation levels in *P. falciparum* (Bridgford et al., 2018; Zhang et al., 2017). Intriguingly, we found that D123 protein levels were reduced by about half after treatment with either Salubrinal or DHA (Fig. 3e-f), opposing the increased transcript levels observed after DHA treatment (Fig. 3a-c). Such an initial decrease of protein levels after activation of the ISR could be explained by the rate at which the mRNA of interest is translated: It is conceivable that protein levels of mRNAs with high ribosome occupancy drop more dramatically relative to the overall proteome, when translation is arrested globally after ISR activation. When we re-analyzed previously published polysome profiling data of ring stage parasites (Bunnik et al., 2013), we found that D123 is a transcript with high polysome occupancy, supporting this hypothesis (log2(polysomal fraction TPM / steady state fraction TPM) = 2.59, compared to median of 2.04 for the overall *P. falciparum* transcriptome) (Fig. 3g, left). We reasoned that other markers of the damage response in cluster 1 might also represent highly translated transcripts. Indeed, cluster 1 markers were highly enriched for transcripts with a high polysome occupancy (log2(polysomal fraction TPM / steady state fraction TPM) = 3.08, compared to median of 2.04 for the overall *P. falciparum* transcriptome, p = 1.24e-5) (Fig. 3g, right).

We were curious whether K13^R539T^ early rings display an analogous pattern of D123 expression, with reduced D123 protein levels contrasting the increased subpopulation of parasites characterized by high D123 transcript levels we had observed (Fig. 1). Indeed, D123 protein levels turned out to be reduced by about 50% at the resistance-relevant early ring stage (0-6 hpi) in the Kelch13 mutant context (Fig. 3h-i).

Taken together, these data show that D123 expression levels are responsive to stress, and that the changes induced by activation of the ISR mimic those observed for untreated early K13-mutant rings, with an expansion of a subpopulation marked by high D123 transcript levels but reduced D123 protein levels. This suggests that the pre-existing activation of the ISR in K13-mutant early rings (Zhang et al., 2017) likely is responsible for the changes in D123 expression we observed at that stage (Fig. 1, Fig. 3h-i).

#### PfD123 expression levels modulate Artemisinin sensitivity

Since D123 proteins positively regulate protein translation, we hypothesized that the pre-existing, reduced D123 protein levels observed in Kelch13-mutant early ring stage parasites might promote resistance to artemisinin. In order to test this, we knocked down D123 protein levels using the glmS ribozyme (Fig. 4a-b), which had no significant effect on asexual growth rates (Supplementary Fig. 6a).

**Figure 4:**
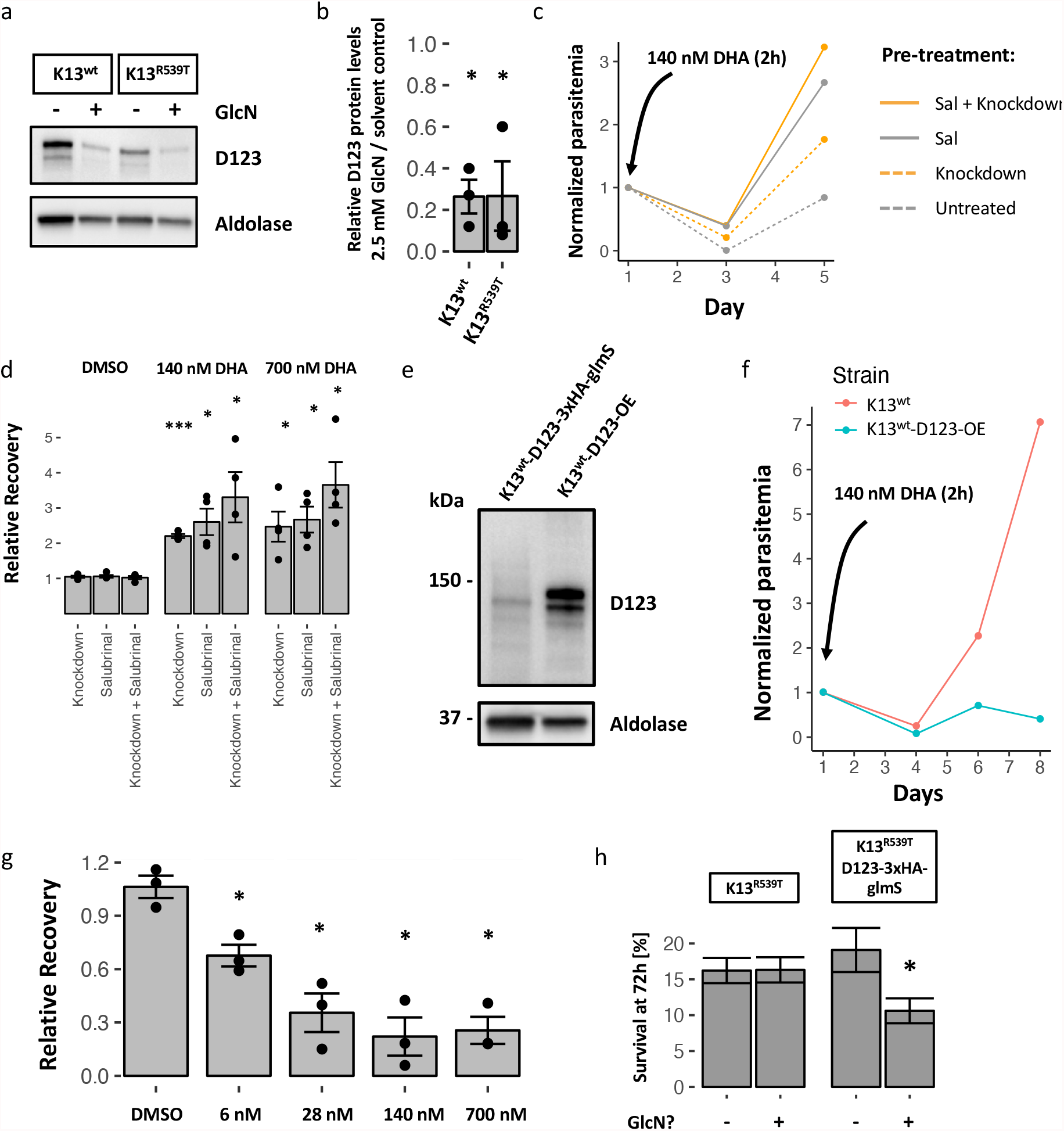
PfD123 expression levels modulate artemisinin sensitivity. **a**, representative immunoblot of D123 levels in K13^wt^-D123-3xHA-glmS and K13^R539T^-D123-3xHA-glmS parasites after 48h treatment with either 2.5 mM glucosamine or solvent control. **b**, densitometry-based quantification of D123 protein levels after 48h knockdown using 2.5 mM glucosamine, plotted as the ratio of Aldolase-normalized D123 expression in glucosamine-over solvent-control treated parasites, n = 3, * = p < 0.05. Representative replicates shown in Fig. 4a. **c**, representative time-course of K13^wt^-D123-3xHA-glmS parasites after different pre-treatments (see figure legend, and methods chapter 3.4.3) and subsequent 2h pulse with 140 nM DHA. Parasitemia is normalized to the starting parasitemia of the individual treatment condition. **d**, quantification of recovery after pre-treatment and DHA pulse (0 / 140 / 700 nM). Relative recovery is defined as the normalized parasitemia at the end of the experiment, relative to parasites that did not receive any pre-treatment. n = 4, * = p < 0.05, *** = p < 0.001. **e**, representative immunoblot of D123 levels in K13^wt^-D123-3xHA-glmS and K13^wt^-D123-OE parasites. **f**, representative time-course of K13^wt^ and K13^wt^-D123-OE parasites after a 2h pulse with 140 nM DHA. Parasitemia is normalized to the starting parasitemia of the individual treatment condition. **g**, quantification of recovery of K13^wt^-D123-OE parasites after DHA pulse (0 / 6 / 28 / 140 / 700 nM). Relative recovery is defined as the normalized parasitemia at the end of the experiment, relative to Dd2 wildtype parasites; n = 3, * = p < 0.05. **h**, percent survival in ring survival assays, of K13^R539T^ and K13^R539T^-D123-3xHA-glmS parasites treated with 0 mM or 2.5 mM glucosamine throughout the whole 72h experiment. n = 5, * = p < 0.05. Error bars always represent sem, statistical significance in b, d and g was tested using two-sided, one-sample t-tests (mu = 1); in h it was calculated using a two-sided, unpaired t-test. In a and e, D123 was detected using an anti-HA antibody, and PfAldolase was used as a loading control. Sal = Salubrinal, DHA = Dihydroartemisinin. GlcN = glucosamine.

However, knockdown of D123 prior to artemisinin treatment in K13^wt^-3xHA-mScarlet-glmS parasites promotes survival after artemisinin exposure at rates similar to pre-treatment with Salubrinal (Fig. 4c-d), treatment with which had previously been demonstrated to promote ART-resistance (Zhang et al., 2017). This suggests that reduced D123 protein levels protect from artemisinin action, potentially by helping with shutting down translation more efficiently at the moment of drug exposure. We reasoned that increased D123 expression should conversely promote sensitivity to artemisinin. As expected, parasites that overexpressed D123 (Fig. 4e, Supplementary Fig. 6b-c) showed delayed recovery after artemisinin treatment (Fig. 4f-g). While reduced D123 levels at the moment of artemisinin exposure are beneficial, we hypothesized that after treatment, when parasites recover from the stress and need to reinitiate protein translation, they require D123 expression at high levels. To test this, we performed RSAs with K13^R539T^-3xHA-glmS parasites, with or without glmS-mediated knockdown of D123 throughout the whole 72h experiment. Indeed, continuous knockdown of D123 reduced survival of K13-mutant parasites (Fig. 4h), suggesting that expression of D123 is critical during later stages of recovery from DHA-inflicted cellular damage.

## DISCUSSION

In this study, we employed scRNAseq to examine differences in the transcriptional heterogeneity between wildtype and artemisinin-resistant malaria parasites at the resistance-relevant early ring stage, and to deconvolute the transcriptional response of parasites to artemisinin treatment. We characterize one gene in more detail: The putative translational regulator D123 that is differentially expressed in K13-mutant early ring stage parasites, with expansion of a subpopulation marked by high D123 transcript levels, but contrastingly reduced D123 protein levels at the same stage. Analogous changes in D123 expression are observed after experimental activation of the ISR, establishing it as a stress-responsive gene and suggesting that the changes in D123 expression observed in K13-mutant rings is a result of the previously reported increased base-line activation of the ISR in K13-mutant parasites (Mok et al., 2015; Zhang et al., 2017). Lastly, by genetic modulation of D123 expression levels, we show that low D123 expression at the moment of artemisinin exposure promote survival, while higher expression levels are needed later, when parasites attempt to recover from the stress.

The integrated stress response is poorly characterized in *Plasmodium* parasites, but it shows great promise as a drug target (Dogovski et al., 2015; Kirkman et al., 2018; Bridgford et al., 2018). Here, we provide new insights into how parasites respond to cytotoxic stress transcriptionally. Our data highlight how heterogeneous the response to stress is, with two transcriptionally different clusters forming in response to DHA treatment, a distinction that has remain masked when bulk transcriptomics approaches have been applied previously (Natalang et al., 2008; Hu et al., 2010; Shaw et al., 2015; Rocamora et al., 2018). The immediate stress response (cluster 2, Fig. 2e top panel) includes Hsp70-1, which has been previously shown to be transcriptionally induced in response to amino acid starvation (Pavlovic Djuranovic et al., 2020) and to be induced by the ApiAP2 transcription factor PfAP2-HS (PF3D7_1342900) after heat shock (Tintó-Font et al., 2021), suggesting it may be a central early stress response gene in the *Plasmodium* ISR downstream of various stimuli. Several other chaperones as well as the drug efflux pump MDR1 were also induced. Intriguingly, several proteins with RNA-binding properties were expressed in this cluster, such as the eIF3 subunit eIF3A. The translation initiation factor eIF3 can be part of stress granules (Buchan & Parker, 2009), and eIF3 and some of its subunits have recently been implicated in the specialized translation of subsets of transcripts during development and stress (Lee et al., 2015; Lee et al., 2016), suggesting that eIF3A may serve a similar function in malaria parasites. Expression of the mRNA-decay factor PfCAF1 (Balu et al., 2011) likely regulates abundance of critical mRNAs during stress, and FHA-domains are commonly found in DNA damage and replication stress proteins in other eukaryotes (Durocher & Jackson, 2002). Taken together, this defines a subset of early stress response genes in cluster 2 that are likely bypassing the translational arrest inflicted by the ISR. How this putative translational bypass is regulated in *Plasmodium* parasites remains to be characterized, as there are major differences in their ISR mechanisms, exemplified by the absence of canonical ISR transcription factors such ATF4 and ATF6 (Chaubey et al., 2014).

The stress-responsive cluster 1 represents the later stage of recovering from the damage inflicted by DHA, containing DNA damage response factors, nucleolar proteins, transcriptional regulators and signaling proteins, amongst others (Fig. 2e, bottom panel). Of note, one of the genes in this marker list was PI3K, which had previously been implicated in ART-resistance (Mbengue et al., 2015). Intriguingly, this distinct set of genes was highly enriched for transcripts that show high polysome occupancy in untreated ring stage parasites (Bunnik et al., 2013; Fig. 3f). It is tempting to speculate that the protein levels of the markers of this cluster are reduced after ISR activation as the translational arrest affects highly translated proteins more strongly, but that they are functionally critical for recovery and therefore are transcriptionally induced through unknown mechanism once stress levels subside in order to quickly recover their protein levels. Future work will have to unravel the regulatory mechanisms controlling this gene expression network.

Efficient translational shutdown at the time of artemisinin exposure, as well as eventually overcoming it after drug levels subside are crucial for the parasite’s ability to recover from artemisinin treatment. This idea is supported by the observation that inflating eIF2α phosphorylation levels in malaria parasites by exposing them to Salubrinal before artemisinin exposure provides protection, while treating with Salubrinal after artemisinin treatment reduces survival (Zhang et al., 2017). We identified the putative translational regulator D123 as a potential downstream effector of the late phase of the ISR that likely mediates some of the effects that had previously been observed for Salubrinal (Fig. 4). Analogous to Salubrinal treatment, knockdown of D123 prior to DHA exposure provides protection (Fig. 4c-d), while knockdown during and after DHA treatment reduces recovery (Fig. 4h). This points to a stress response expression program that needs to be finely tuned in order to enable survival after ISR activation. Importantly, we find that D123 and other cluster 1 markers are differently expressed in untreated Kelch13-mutant, artemisinin resistant parasites at the resistance-relevant early ring stage. We propose a model, in which the low-level activation of the ISR in Kelch13-mutant early rings (Zhang et al., 2017) leads to reduced protein levels of D123 and other cluster 1 markers, which promotes survival by supporting an efficient translational shutdown, while the concomitant increased transcript levels resulting from the damage response support recovery once drug levels subside. Importantly, we find that increased D123 transcript levels in response to stress are restricted to a subpopulation of parasites (clusters 1 in Fig. 1a and Fig. 2a). Transient changes in expression of stress-responsive genes that provide protection against artemisinin in some parasites but not others would provide an explanation why it is only ever a subpopulation of Kelch13-mutant parasites that survive exposure to artemisinin, but future work will have to conclusively demonstrate this to be true.

Of note, the D123 Dd2 ortholog PfDd2_030027500, as opposed to its 3D7 ortholog PF3D7_0322400, is currently misannotated as a pseudogene due to an additional Adenine detected within a polyadenine stretch in its open reading frame, which likely is a result of polymerase slippage during reference genome generation (PlasmoDB.org). When we tagged the endogenous PfDd2_030027500 locus, a protein of appropriate size was detectable by immunoblot, and when amplifying PfDd2_030027500 from gDNA, the additional adenine was not present, together clearly demonstrating that this is a protein coding gene. Notably, in K13^wt^-D123-3xHA-mScarlet-glmS and K13^R539T^-D123-3xHA-mScarlet-glmS parasites (Supplementary Fig. 2), where the endogenous D123 locus was tagged both with a 3xHA and a fluorescent mScarlet tag, we were never able to detect mScarlet fluorescence, while the protein was readily detectible by immunoblot using an anti-HA antibody. We confirmed that there were no mutations in the mScarlet tag (data not shown) and hypothesize that fluorescence levels might be too low or potentially too transient, as this protein appears to be highly translated and might only be individually present for a short period of time. Its suggested high rate of translation is supported by the observation that the destabilization domain used in the D123 overexpression line fails to reduce D123 protein levels, leading to high D123 expression even in the absence of Aquashield in the culture media (Supplementary Fig. 3c). Furthermore, when detecting D123 by immunoblotting, we consistently observed two protein bands of similar size, indicating processing of this protein (e.g. Fig. 1f).

Taken together, this work provides important new insights into how *P. falciparum* responds to stress, and into how Kelch13-mutant parasites hijack those mechanisms to gain protection against artemisinin.

## MATERIALS & METHODS

### Parasites and strains

The strains of parasite used in this study were Dd2 (K13^wt^), Dd2^R539T^ (K13^R539T^) (Straimer et al., 2015), as well as additional parasite strains generated during this research project (Supplementary Fig. 2, Supplementary Fig 6, see details below). They were maintained using established cell culture techniques (Moll et al., 2008). K13^wt^ and K13^R539T^ strains were generously gifted by Laura Kirkman and tested for the engineered phenotype, showing expected levels of survival when tested in ring survival assays in June 2019.

### Ring survival assays

Ring survival assays were performed as previously described (Witkowski et al., 2013). Briefly, parasites were tightly synchronized to a 0-3 hpi window using percoll-sorbitol enriched schizonts incubated with fresh red blood cells for 3 hours, after which the remaining schizonts were removed using a 5% sorbitol wash and three washes with incomplete media. 0-3 hpi rings were then exposed to 700 nM DHA or DMSO for 6h, after which the drug was washed off and parasites were allowed to recover. Blood smears were collected at 72h after the start of the experiment in order to count viable parasites and determine parasitemia. Dividing the parasitemia of DHA-treated parasites by the parasitemia of DMSO-treated parasites from the same culture determines the RSA survival.

### Other artemisinin sensitivity assays

For experiments determining ART-sensitivity after D123 knockdown, semi-synchronous K13^wt^-D123-3xHA-glmS parasites were incubated with 2.5 mM glucosamine or solvent control for 24h to mediate knockdown, starting at around 24 hpi. When parasites were invading new red blood cells at around 48 hpi, remaining schizonts were removed using a 5% sorbitol wash. Early rings remained in culture, and were treated with solvent controls, 2.5 mM glucosamine, 10 μM Salubrinal, or both, for 2h. After washing off pre-treatments, parasites were then exposed to 0.1% DMSO, 140 nM DHA or 700 nM DHA for 2h, after which drugs were washed off, and blood smears were used to determine the starting parasitemia for each condition. Recovery was followed up with daily blood smears, which were used to track parasitemia over time. Experiments were terminated when at least one treatment condition grew back beyond its initial starting parasitemia.

Experiments determining ART-sensitivity in K13^wt^-D123-3xHA-OE parasites were set up similarly: Semi-synchronous parasites were incubated with 5% sorbitol when they were reinvading new red bloods cells in order to remove remaining schizonts and recover early ring stage parasites, which were then incubated for 2h with different concentrations of DHA or 0.1% DMSO. Drugs were washed off and blood smears were used to determine the starting parasitemia for each condition. Recovery was followed up with daily blood smears, which were used to track parasitemia over time. Experiments were terminated when at least one treatment condition grew back beyond its initial starting parasitemia.

### Parasite isolation for single-cell RNA-seq

K13^wt^ wildtype and K13^R539T^ parasites were set up for ring survival assays as described above. For the 0-3 hpi experiments (Fig. 1), parasites were processed for Drop-seq immediately after the sorbitol treatment and subsequent three washes with incomplete media used to remove remaining schizonts after the three-hour incubation with fresh uRBCs (see below, ring survival assays). For the RSA 6-9 hpi and 15-18 hpi samples (Fig. 2), parasites were processed for Drop-seq immediately after washing off DHA after the 6h incubation (6-9 hpi), and then again after parasites had been back in culture for an additional 9h after DHA exposure (15-18 hpi), meaning that the two RSA scRNA-seq replicates each were true time-course experiments with both time-point collected from the same culture flask. Parasite processing at the different time-points started with the Streptolysin O-Percoll (SLOPE) method (Brown et al., 2020) to enrich for ring-infected RBCs: the pellet of infected RBCs was incubated with 30 U Streptolysin-O (SLO) per 1e8 RBCs at RT for 6 min, then washed three-times with PBS, and fractionated on a 60% percoll gradient. A small portion of the resulting iRBC-enriched pellet was dried on a glass slide and GIEMSA stained to determine the degree of enrichment, while the rest was resuspended in ice-cold PBS + 0.01% BSA and immediately processed for Drop-seq. For all scRNA-seq experiments, a small amount of culture was not processed and instead kept in culture to confirm expected levels of survival for the different strains by RSA.

### Drop-seq and sequencing analysis pipeline

Single-cell transcriptomic profiles were generated using Drop-seq, a technology designed for highly parallel genome-wide expression profiling of individual cells using nanoliter droplets (Macosko et al., 2015), and applied to malaria parasites as previously described (Poran et al., 2017). In brief, single-cell suspensions and uniquely barcoded beads were colocalized in droplets using a microfluidics device (see CAD file from http://mccarrolllab.com/dropseq/, anufactured by FlowJEM). The droplets are composed of cell-lysis buffer and serve as compartmentalizing chambers for RNA capture. For some of the samples in this study, the cell-lysis buffer was modified to contain 4 M Guanidine-HCl for additional RNase inhibition in addition to the published recipe (Macosko et al., 2015). Flow rates were adjusted to maintain stable droplet formation and increase droplet homogeneity, yielding 1 nanoliter droplets in our setup. For some of the samples in this study, the oil flow rate was increased to reduce droplet size with the goal to improve RNA capture efficiency by increasing the local RNA concentration within the droplet during lysis. The maximum oil flow rate that still allowed for stable droplet formation and homogeneous droplets yielded 500 picoliter droplets. For each setting (1 nl versus 500 pl), cell and bead concentrations were adjusted to accommodate variation in droplet size compared to the original publication (Macosko et al., 2015).

Droplet breakage and single-cell library preparations followed the procedure as described (Macosko et al., 2015). In brief, collected droplets were disrupted and RNA-hybridized beads were extracted. Reverse transcription was performed with template switching to allow for cDNA amplification by PCR. An additional pre-PCR step was added to determine the appropriate number of cycles (31-33 cycles for 0-3 hpi samples, 27-28 cycles for 6-9 hpi samples, 24-27 cycles for 15-18 hpi samples) to achieve a cDNA library at a concentration of 400–1,000 μg μl^−1^, as suggested by the protocol. cDNA samples were purified using Agencourt AMpure XP (Beckman Coulter), and were run on a 2100 BioAnalazyer instrument with a High Sensitivity DNA kit (Agilent Technologies). Samples were prepared for sequencing using the Illumina Nextera XT kit, and sequenced on a NextSeq 500 (Illumina) at an average of 50,000 reads per cell. Raw reads were processed and aligned (STAR aligner) using the standard Drop-seq pipeline, and according to the ‘Drop-seq Alignment Cookbook’, both found at http://mccarrolllab.com/dropseq/. Reads were aligned to the polyadenylated *P. falciparum* Dd2 transcriptome (PlasmoDB v. 43). For each read, a single optimal mapping position was retained. Unique transcripts mapping to alternative splice variants were combined for subsequent analysis. Single-cell expression matrices were generated using cellular barcodes and unique molecular identifiers (UMIs). Transcript capture from the different parasite stages improved progressively with increasing hpi of the samples, reflecting the increasing RNA content of parasites as they progress through intra-erythrocytic development (Martin et al., 2005, Supplementary Fig. 1b, Supplementary Fig. 3b).

### Single-cell transcriptome analysis

Data normalization, clustering and differential expression were performed using the Seurat R package (Satija et al., 2015). Cells with less than 10 (0-3 hpi replicates 1 and 3), 5 (0-3 hpi replicate 2) or 50 (6-9 and 15-18 hpi) UMIs and genes detected in fewer than three cells were excluded from analysis. SCTs were internally normalized to 10,000 transcripts, log transformed and regressed on the number of UMIs per cell before dimensionality reduction and clustering. We selected ∼600–700 highly variable genes using the expression and dispersion (variance/mean) of genes, and performed principle component analysis. The most significant principal components (heuristically determined based on PC significance elbow plot) were used for clustering and UMAP representations (Becht et al., 2018). Clustering resolution was chosen such that visually distinct groups of cells were assigned to individual clusters.

### SCT correlation with bulk RNA-seq data

Mapping the clusters 3–5 of the RSA scRNAseq experiments to the time series bulk RNA-seq dataset (Bártfai et al., 2010) was performed by pseudo-bulking the SCTs within each cluster and then calculating the Pearson’s correlation coefficient of the clusters with each of the 8 bulk-RNA-seq time points. The two scRNAseq replicates were analyzed separately.

### Plasmid constructions and generation of novel engineered parasite strains

K13^wt^-D123-3xHA-mScarlet-glmS and K13^R539T^-D123-3xHA-mScarlet-glmS parasite lines (also referred to as K13^wt^-D123-3xHA-glmS and K13^R539T^-D123-3xHA-glmS for brevity) were generated by co-transfecting K13^wt^ and K13^R539T^ respectively with pUF1-Cas9-yDHDOH and pD123-3xHA-mScarlet-glmS (Supplementary Fig. 2a), and selecting with DSM1 and G418 until viable parasites were recovered by blood smear. Successful editing of the genomic D123 locus was verified by PCR using a forward primer outside of the 5’ homology block and reverse primers binding in the recoded 3’ end of the gene or in the mScarlet tag (Supplementary Fig. 2b). Constructs were designed such that the single intron of PfDd2_030027500 as well as a poly-adenine stretch just 3’ of the stop codon were removed in the transgenic lines. Since the genomic locus was tagged without integration of selection marker expression cassettes, drug selection was dropped once correct editing had been verified.

K13^wt^-ddFKBP-mScarlet-3xHA-D123 parasites (also occasionally referred to as K13^wt^-D123-OE for brevity) were overexpressing tagged D123 using a calmodulin promoter, and were generated by transfecting pddFKBP-mScarlet-3xHA-D123-OE into K13^wt^ parasites (Supplementary Fig. 6b) and selecting with Blasticidin S until viable parasites were recovered by blood smear. Successful overexpression was verified by immunoblot. Blasticidin S selection was maintained for continuous culture, but was dropped at the beginning of experiments.

### Protein extractions and immunoblotting

Parasites were released from red bloods by incubating RBCs with 0.05% saponin in PBS for 10 min on ice. The pellet was washed with PBS until the supernatant showed no more signs of hemoglobin release. Proteins were isolated from parasite pellets using MPER lysis buffer (Thermo Fisher) supplemented with 1x c0mplete protease inhibitors (Sigma-Aldrich). A small aliquot was set aside for protein quantification using BioRad Protein Assay, while the rest was supplemented with SDS-loading buffer and boiled at 95C for 5 min. Pellets were then frozen and stored at -80C until use.

10 μg of protein extracts were separated on a 4-20% gradient SDS-PAGE TGX stain-free gels and transferred to PVDF membranes using an iBlot (transfer program “P0”). Total protein was visualized on membranes using BioRad stain-free crosslinking technology. Membranes were then blocked with 5% milk powder in TBS-T (0.1% tween), and incubated with primary antibodies for 1h at RT or at 4C overnight (1:2,500 rat anti-HA; 1:5,000 rabbit anti-aldolase; in 5% milk powder in TBS-T). After three washes with TBS-T, secondary antibodies were incubated for 1h at RT (1:10,000 rabbit HRP-anti-rat; 1:10,000 goat HRP-anti-rabbit; in 5% milk powder in TBS-T). After three more washes in TBS-T chemiluminescence-based visualization was performed using SuperSignal West Pico PLUS substrate (Thermo Fisher) or SuperSignal West Femto Maximum Sensitivity substrate (Thermo Fisher). Densitometry-based quantification of immunoblot signals was performed using ImageJ imaging software.

### Analysis of previously published polysome profiling data

TPM counts of whole-transcriptome analyses of steady-state mRNA fractions and polysome-associated mRNA fractions collected at the ring stage (Bunnik et al., 2013) were downloaded from PlasmoDB.org. Only uniquely mapped reads were analyzed. Genes for which no steady-state or polysome-associated reads were detected were removed from the dataset. Polysome-occupancy was estimated for each gene by calculating the log2 of the ratio (polysome-associated TPM / steady-state TPM).

### Other statistical analyses

Significance of cluster marker analysis was assessed using the Wilcoxon Rank Sum test, *P* values in multiple hypothesis testing are FDR corrected (Bonferroni correction). No statistical methods were used to predetermine sample size. The experiments were not randomized. The investigators were not blinded to allocation during experiments and outcome assessment.

## General Acknowledgements

We would like to thank the WCM Genomics core facility, as well as Olivier Elemento for generously gifting the Drop-seq setup to the Kafsack laboratory, and Laura Kirkman for gifting the K13^wt^ and K13^R539T^ parasite lines. This work was supported by WCM internal startup funds (B.F.C.K.) and a US Department of Defense CDMRP PRMRP-Discovery Award W81XWH-18-1-0222 (B.F.C.K.). C.N. was supported by WCM graduate fellowships and the Jacques Cohenca Predoctoral Fellowship.

## Contributions

B.F.C.K. conceived the study and wrote a pearl script for polyadenylated reference transcriptome generation. C.N. performed all experiments, analyzed the data, designed and generated the figures and wrote the manuscript.

## SUPPLEMENTARY FIGURES

**Supplementary Figure 1:**
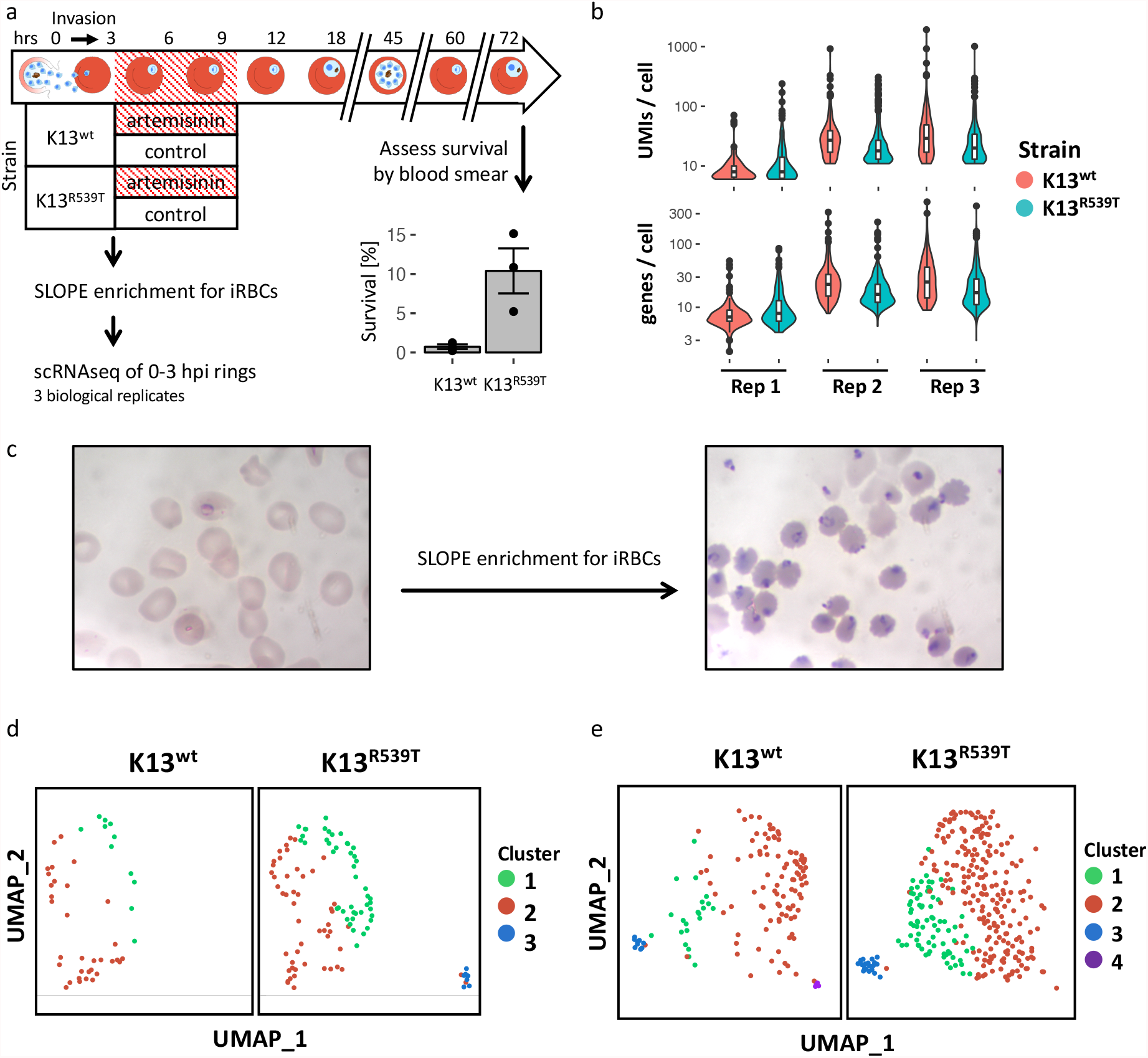
scRNAseq of artemisinin-sensitive and -resistant 0-3 hpi ring stage parasites. **a**, experimental setup for early ring scRNAseq experiments and corresponding RSA survival data. Tightly synchronous 0-3hpi ring parasites were enriched for infected red blood cells using the SLOPE method (Brown et al., 2020), then immediately processed for scRNAseq. Some remaining culture was kept aside and followed up to determine relative survival rates (bar plot in bottom right corner, n = 3, p = 0.076, two-sided unpaired t-test, error bars represent sem). **b**, unique molecular identifiers (UMIs, top) and genes (bottom) detected per cell in the six 0-3 hpi scRNAseq samples. **c**, representative Giemsa smears of 18 hpi ring stage parasites before (left) and after (right) enrichment for iRBCs using SLOPE. **d, e**, UMAP projections of replicate 2 (d) and replicate 3 (e) of the 0-3 hpi scRNAseq experiments, stratified by parasite strain.

**Supplementary Figure 2:**
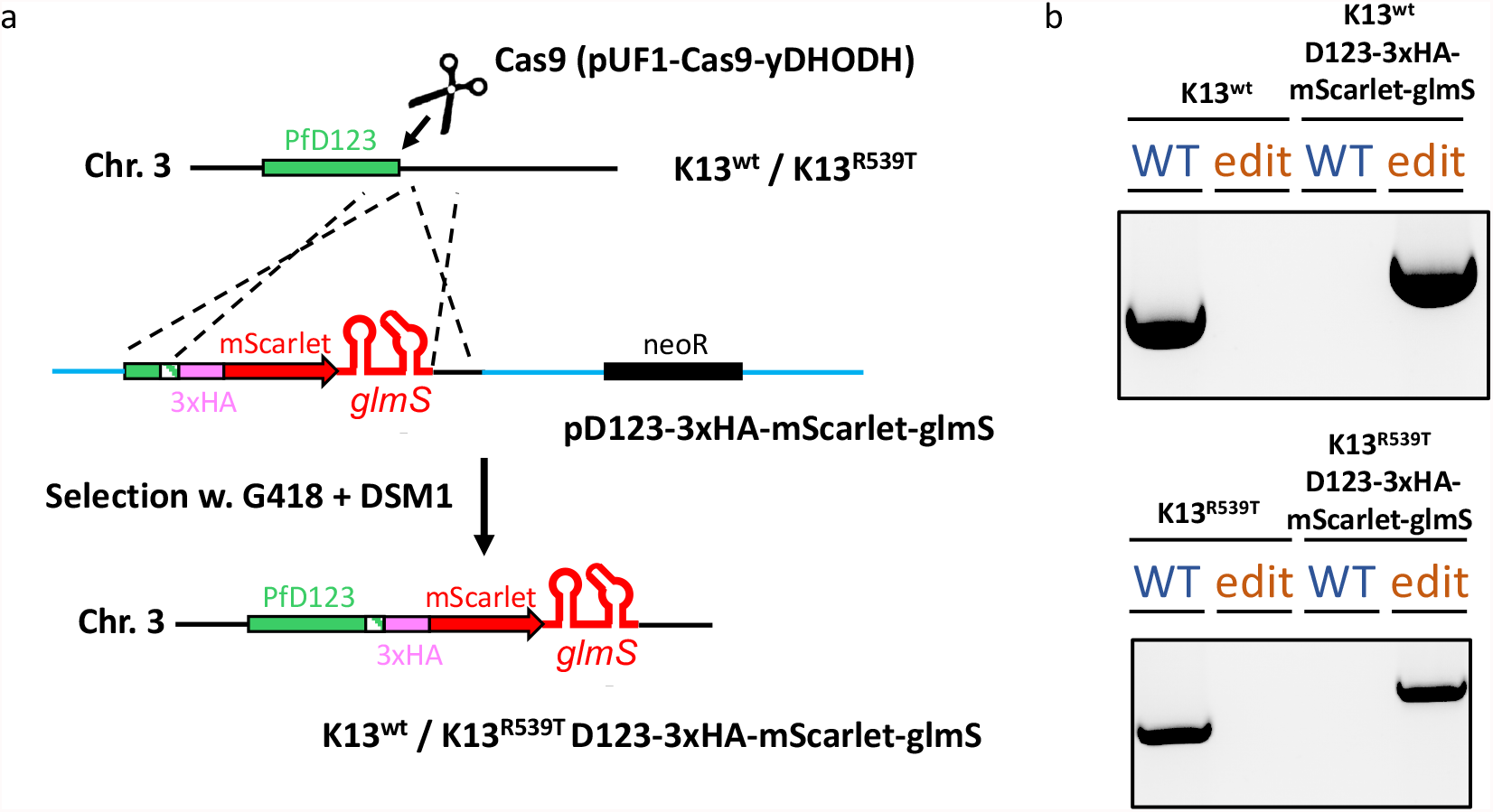
Generation of epitope-tagged D123 knockdown parasite lines in K13^wt^ and K13^R539T^. **a**, overview of the strategy for generating K13^wt^-D123-3xHA-mScarlet-glmS and K13^R539T^ -D123-3xHA-mScarlet-glmS parasite lines. Green striped area indicates part of the ORF between Cas9 cut-site and STOP codon that was recoded. **b**, PCR validation of successful gene editing. “WT” and “edit” PCRs both share same forward primer positioned within the ORF, outside the 5’ homology region. Reverse primer for “WT” PCR was positioned in sequence that ended up recoded after editing, while reverse primer for “edit” PCR was positioned within the mScarlet tag.

**Supplementary Figure 3:**
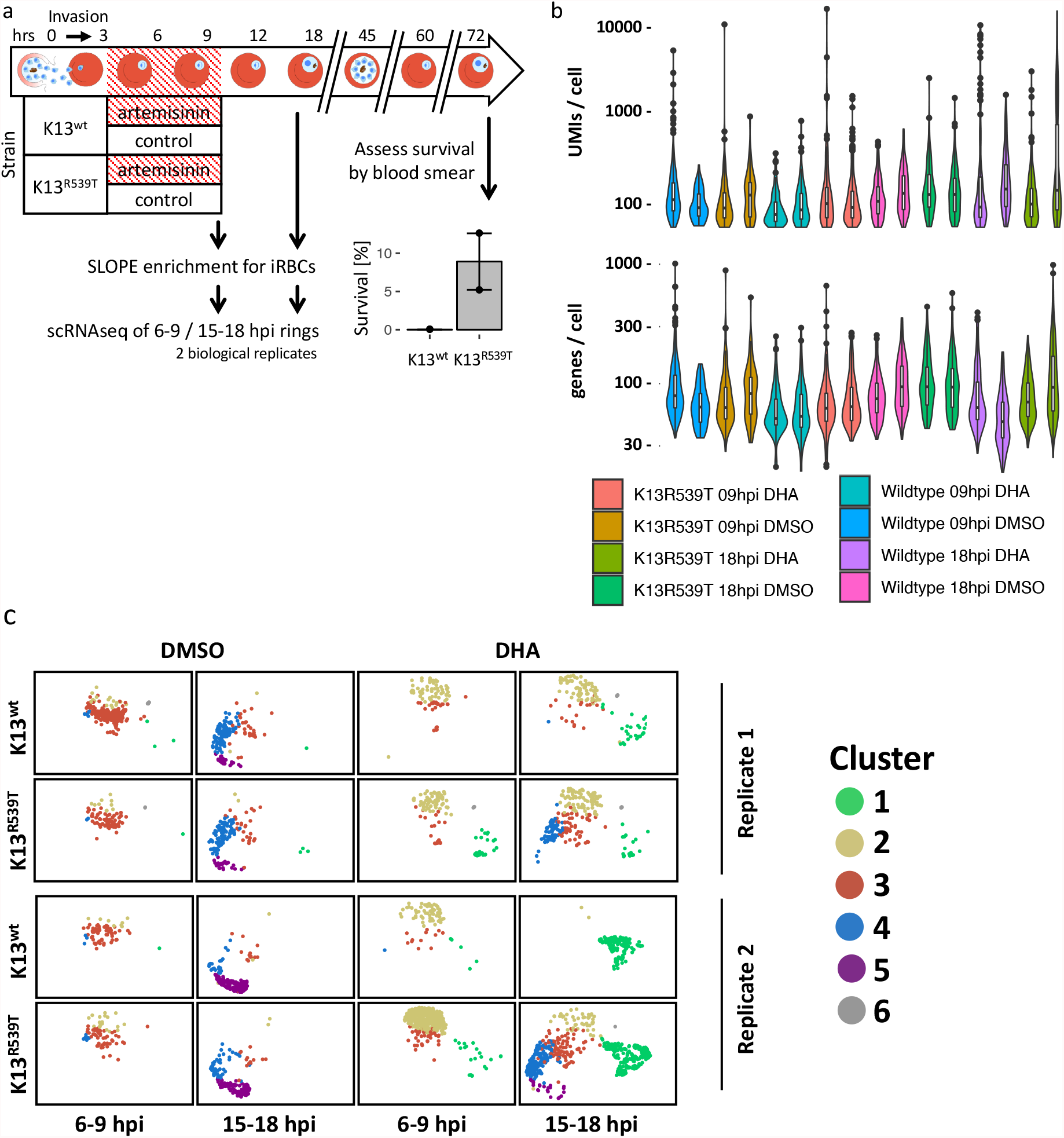
scRNAseq of artemisinin-sensitive and -resistant parasites after artemisinin treatment. **a**, experimental setup for scRNAseq experiments of ring stage parasites after artemisinin treatment and corresponding RSA survival data. Tightly synchronous 0-3hpi ring parasites were exposed to 0.1% DMSO or 700 nM DHA, and samples were collected at 6-9 hpi and 15-18 hpi, respectively. Parasites were enriched for infected red blood cells using the SLOPE method (Brown et al., 2020), then immediately processed for scRNAseq. Some remaining culture was kept aside and followed up to determine relative survival rates (bar plot in bottom right corner, n = 2). **b**, unique molecular identifiers (UMIs, top) and genes (bottom) detected per cell in the experiments described in a. **c**, UMAP projections all experiments described in a, stratified by parasite strain, collection time point, treatment and biological replicate.

**Supplementary Figure 4:**
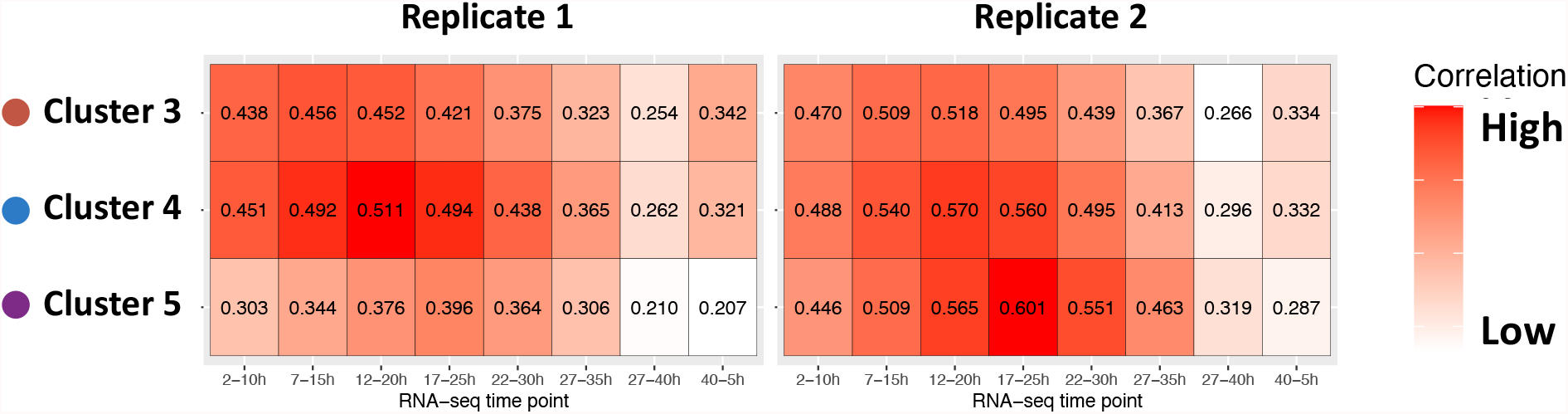
Parasites progress through intra-erythrocytic development from cluster 3 to 5 in the RSA scRNAseq experiments. SCTs of clusters 3-5 were pseudo-bulked and then correlated to the individual time points of a time-series RNA-seq bulk transcriptomics dataset (Bartfai et al., 2010). Tile color shading is based on Pearson’s correlation coefficient of the respective pair-wise comparison, with the coefficient printed on each tile.

**Supplementary Figure 5:**
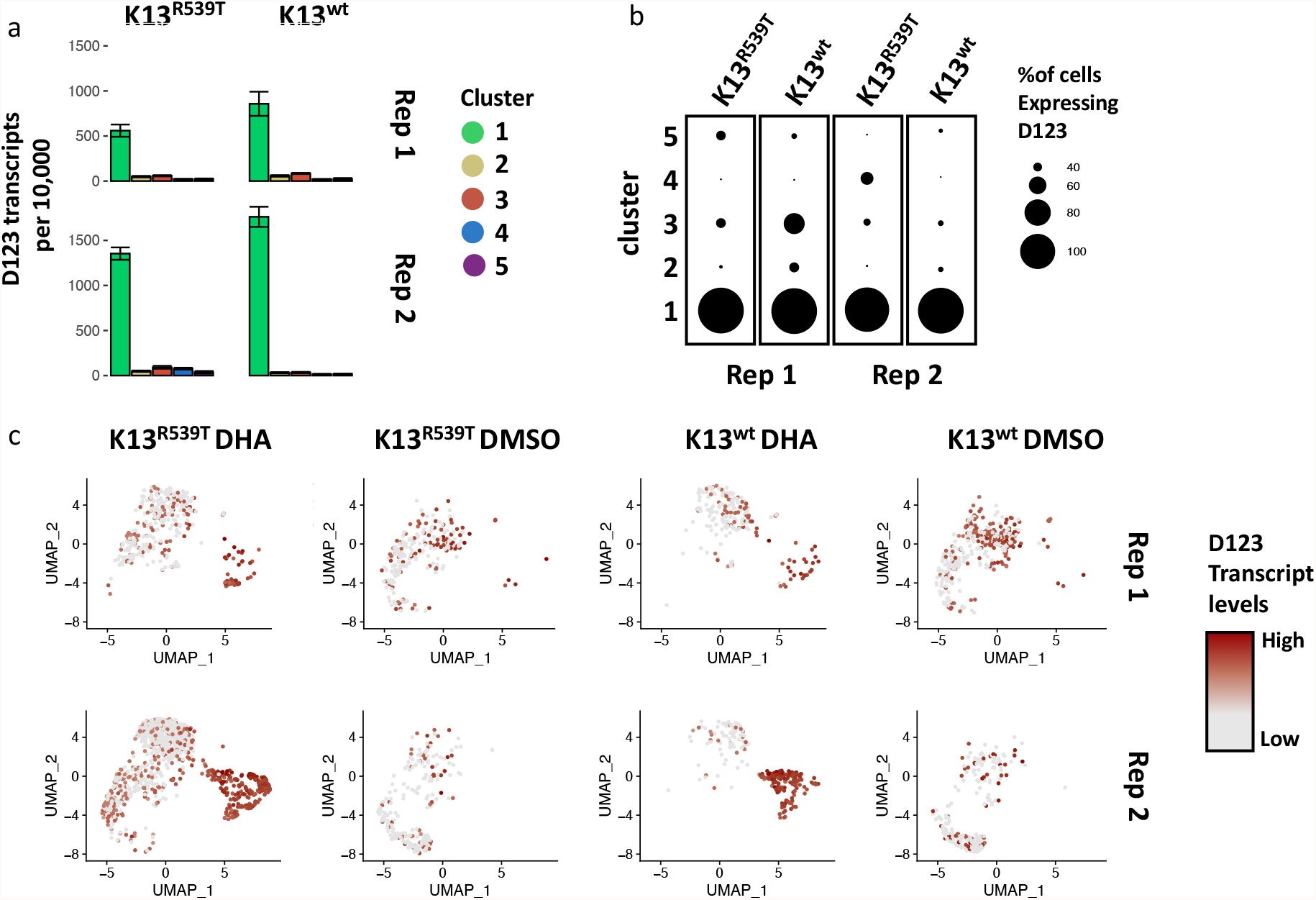
Expression of D123 in response to artemisinin. **a**, same data as in Fig. 3a, but stratified by strain and replicate. Error bars represent sem. **b**, same data as in Fig. 3b, but stratified by strain and replicate. **c**, UMAP projections of all SCTs of the RSA experiments (Fig. 2), stratified by treatment, strain and replicate, with each cell colored based on the detected D123 transcript levels. DHA = Dihydroartemisinin.

**Supplementary Figure 6:**
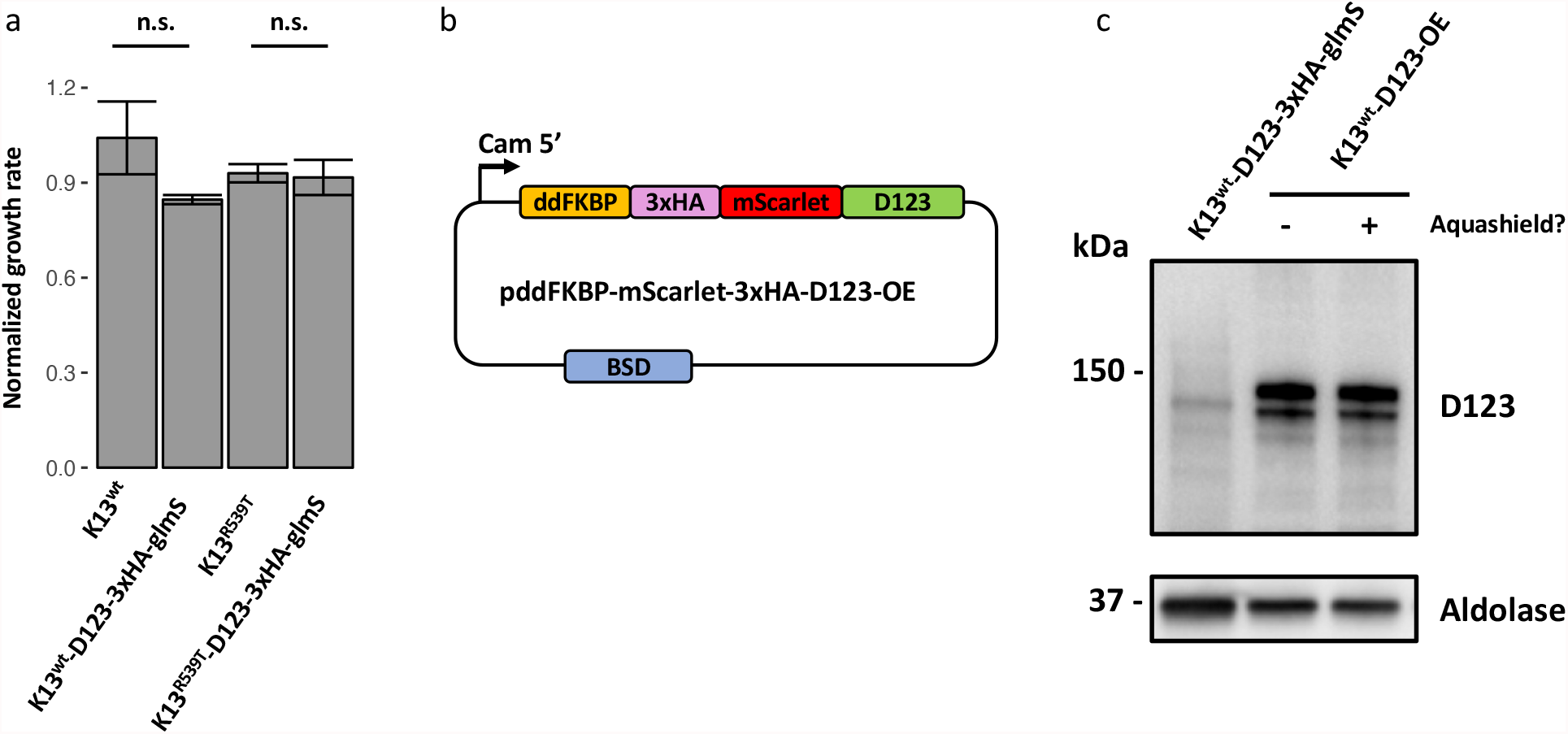
Modulation of D123 expression levels. **a**, growth rates of different parasite strains after 2.5 mM glucosamine treatment, plotted normalized to the growth rate when treated with a solvent control. **b**, schematic representation of the plasmid map and strategy used for overexpression of D123. Similar to the parasite lines where the endogenous D123 locus was tagged, little to no fluorescence was detected for D123-OE parasites (data not shown). **c**, Same representative immunoblot as shown in Fig. 4e, but including an additional lane showing D123 expression levels in K13^wt^-ddFKBP-3xHA-mScarlet-D123-OE parasites after 24h pretreatment with 500 nM Aquashield. ddFKBP is supposed to target D123 to the proteasome, rendering D123 overexpression regulatable, as it should only be stabilized upon addition of Aquashield to the culture media. However, D123 protein levels were high in this line even in the absence of Aquashield in the media. D123 was detected using an anti-HA antibody, and PfAldolase was used as a loading control.

